# Deficiency of cyclin-dependent kinase-like 5 causes spontaneous epileptic seizures in neonatal mice

**DOI:** 10.1101/2020.03.09.983981

**Authors:** Wenlin Liao, Kun-Ze Lee, San-Hua Su, Yuju Luo

## Abstract

Cyclin-dependent kinase-like 5 (*CDKL5*), an X-linked gene encoding a serine-threonine kinase, is enriched in the mammalian forebrain and critical for neuronal maturation and synaptic function. Mutations in this gene cause CDKL5 deficiency disorder (CDD) that is characterized by early-onset epileptic seizures, autistic behaviors and intellectual disability. Although numerous CDD symptoms have been recapitulated in mouse models, spontaneous seizures have not been reported in mice with CDKL5 deficiency. Here, we present the first systematic study of spontaneous seizures in a mouse model of CDD. Through wireless electroencephalographic (EEG) recording and simultaneous videotaping, we observed epileptiform discharges accompanied with ictal behaviors in pups lacking CDKL5 at a selective time window during the pre-weaning period. The seizure-like patterns of EEG showed robust increase in total number of spike events, the total number and duration of bursts in *Cdkl5* null pups compared to wild-type littermate controls at the age of postnatal day 12 (P12). The mutants displayed not only jerky and spasm-like movements during the prolonged bursts of discharges at P12, but also strengthened ictal grasping in both juvenile stage and adulthood. In addition, loss of CDKL5 remarkably reduced the phosphorylation of K^+^/Cl^-^ co-transporter 2, which may impede GABA-mediated inhibition, in the cortex of P12 mouse pups. Our study reveals previously unidentified phenotypes of early-onset seizures in CDKL5-deficient mice, highlights the translational value of mouse models of CDD and provides a potential molecular target for early diagnosis and treatment for CDD.

**Significance Statement:** Cyclin-dependent kinase-like 5 (*CDKL5*) is an X-linked gene encoding a serine-threonine kinase. Mutations in this gene cause CDKL5 deficiency disorder (CDD), a rare disease characterized by developmental delays, autistic behaviors and early-onset epilepsy. Even though many symptoms of CDD patients have been phenocopied in mice, spontaneous seizures are yet to be reported in mouse models of CDD. Here, for the first time, we identified early-onset seizures and ictal behaviors in neonatal pups of CDKL5-deficient mice. Loss of CDKL5 also selectively reduced protein levels of phosphorylated K+/Cl-cotransporter 2 in neonatal cortex of mice. Our study reveals an indispensible role of CDKL5 in regulating neuronal excitability in developing brains and highlights the translational significance of the CDD mouse models.

## Introduction

CDKL5 deficiency disorder (CDD; OMIM#300672; ICD-10-CM code: G40.42) is a developmental and epileptic encephalopathy caused by mutations in the X-linked gene encoding cyclin-dependent kinase-like 5 (CDKL5; OMIM#300203). This disorder is characterized by severe early onset seizures, autistic features, and intellectual disability (Weaving et al., 2004; Scala et al., 2005; Archer et al., 2006), and categorized as one of the most common forms of genetic epilepsy (Lindy et al., 2018; Symonds et al., 2019). The epileptic activities accompanied with developmental delays in children with CDD are frequently associated with devastating cognitive, psychiatric and behavioral disabilities later in life. Elimination of early-onset epilepsy may potentially ameliorate the consequent developmental delays of this disorder (Fehr et al., 2016; Scheffer et al., 2017; Demarest et al., 2019). Despite well-defined genetic cause, however, children with CDD are resistant to most of the anti-epileptic drugs (AEDs) and the pathogenic mechanisms for CDKL5 deficiency-induced seizures remain unclear.

Several mouse models of CDD have been developed that display behavioral phenotypes mimicking primary symptoms of CDD, but intriguingly, spontaneous seizures have not been reported (Wang et al., 2012; Amendola et al., 2014; Jhang et al., 2017; Okuda et al., 2018). In the study of the first mouse model, normal basal electroencephalographic (EEG) patterns were recorded in the hippocampus of *Cdkl5* null mice carrying a deletion of exon 6 using deep electrodes for up to 240 hours (Wang et al., 2012). Upon kainic acid administration, mice lacking exon 4 of *Cdkl5* gene show epileptiform discharges, but normal basal EEG, in the somatosensory cortex (Amendola et al., 2014). Moreover, spontaneous seizures have not been found in the third CDD mouse model with deletion of *Cdkl5* exon 2, although mutant mice exhibited enhanced seizure susceptibility in response to NMDA treatment (Okuda et al., 2017). Notably, all the behavioral observations and EEG recordings for seizure activities in these studies were conducted in adult mice aged from 1 to 4 months. Considering that the median age of seizure onset in CDD patients is about 6 weeks after birth and a “honeymoon period” with seizure cessation can last up to 72 months (Bahi-Buisson et al., 2008; Fehr et al., 2016; Demarest et al., 2019), it is possible that early-onset spontaneous seizures were missed in these studies of adult mice.

We previously noticed that *Cdkl5* null (exon 6 deletion, *Cdkl5*^*-/y*^) pups showed increased spasm-like twists compared to wild-type (WT) pups during recordings for ultrasound vocalization at the early postnatal period (P4 - P10) (Jhang et al., 2017)(Liao, unpublished observations). The similar spasm-like behaviors have been described in mice with *Arx* gene mutations that cause infantile spasm (Marsh et al., 2009; Price et al., 2009), suggesting that pups lacking CDKL5 may experience early-onset epilepsy soon after birth. Given that EEG recordings and seizure behaviors have not been reported in pre-weaning pups with CDKL5 deficiency, in this study, we thus carried out longitudinal EEG recordings coupled with simultaneous videotaping in mouse pups. To reveal possible molecular targets for therapeutic development, we also examined the expression of candidate proteins affected by CDKL5 deficiency in neonatal mice.

## Materials and Methods

### Animals

Male *Cdkl5* null pups were generated by crossing C57BL/6J male mice (National Laboratory Animal Center, Taiwan) to heterozygous *Cdkl5* females [*Cdkl5*^+*/-*^, B6.129(FVB)-*Cdkl5*^*tm1*.*1Joez*^/J](Wang et al., 2012). Male hemizygous pups (*Cdkl5*^*-/y*^) and their littermate controls (*Cdkl5*^+*/y*^) were used as the experimental cohorts to avoid mosaic expression of *Cdkl5* due to random X-inactivation. All mice were bred and housed in individually ventilated cages (IVC, Alternative Design, USA) at 22 ± 2°C and 60 ± 10 % humidity under a 12-hour light-dark cycle (light on 08:00 to 20:00). Irradiated diet (#5058, LabDiet, USA) and sterile water were supplied *ad libitum*. All experiments were approved by the Institutional Animal Care and Use Committee at National Cheng-Chi University.

### Pup EEG recording - Implantation of transmitters

Male pups were tail-marked and genotyped at P9. Genotyping was performed as previously described (Jhang et al., 2017). Only the pups with a body weight of > 6 g at P10 were subjected to transmitter implantation. Isoflurane anesthesia (1% in O_2_) was applied to each pup for ∼20 min during the surgery. The transmitters (4 × 6 mm in size of footprint, 8 mm in height, 0.5 g for each; gain = 4000 x) were activated before use and have a two-week guarantee according to the manufacturer’s instructions (Epoch, Epitel, USA) (Zayachkivsky et al., 2013). The bilateral electrodes (2 channels, ∼2 mm in between) on the activated transmitters were cut to ∼2-3 mm and then implanted at the middle level of the mouse cortex symmetrically to the midline, and a reference electrode (∼3 mm) was positioned to the left anterior to the Lambda. After the transmitter was secured onto the skull with tissue glue (Vetbond, 3M), the implanted pups were kept on a pre-warmed plate until awakening and then monitored for EEG activity for 20 min to confirm success of implantation (Fig. 1). The implanted pups were then returned to their home cage and monitored for their body weight and general health on a daily basis to confirm their recovery and growth. Six pairs of pups were successfully implanted and survived. Due to unexpected failure of the transmitters during the experiments (P11-P24), however, only four pairs (from four litters) of data were adopted for statistic analysis.

**Figure 1.**
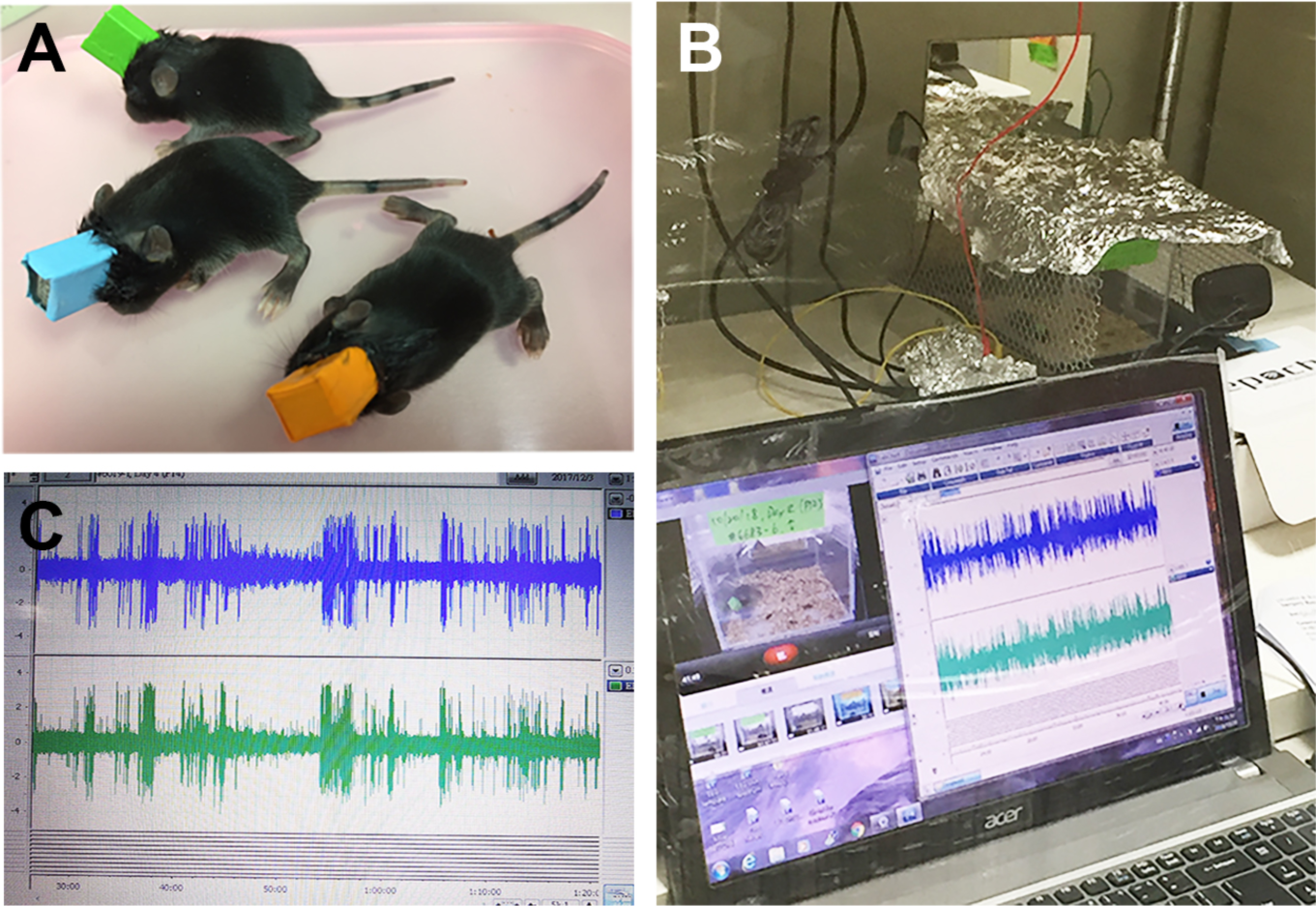
Wireless EEG recordings performed in mouse pups. A transmitter, including electrodes, a battery and an antenna in an epoxy-coated cube of 4 × 6 mm^3^ in size was implanted in a mouse pup at postnatal day 10 (P10, A). The implanted pup was placed in a transparent acrylic box on the top of the receiver and subjected to synchronized recording of EEG and behaviors (B). Seizure-like EEG activities were acquired at the sampling rate of 1000 Hz (C).

### Pup EEG recording - Data acquisition and analysis

For longitudinal EEG studies, these pups were placed in a transparent plastic box (10 cm x 10 cm) and recorded daily between 1-6 pm for 100 min in the first four days post-surgery (P11-14), followed by recording at P17, 21, and 24. EEG activities were captured with a receiver (Epoch, Epital) connected to a data acquisition system (PowerLab, ADInstruments) controlled by the LabChart 7 software. The sampling rate was 1000 Hz. The raw data were first processed by digital filter with a cut-off frequency at 50 Hz (low-pass) and 3 Hz (high-pass), and then the traces for 10-100 min (90 min) were exported to Clampex (Version 10.7, Axon Instruments). For event detection, the threshold was set as 0.9 V (equivalent to 0.225 mV of input intensity) to detect spike discharges (i.e. events; Fig. 4A). For the burst analysis, a burst was defined as a continuous train of discharges that contains more than 20 spikes with the inter-spike interval of < 1 sec. The measurements for event/burst parameters (duration, frequency, and amplitude) were exported to Excel for statistic analysis (Table 4-1). Given that the EEG data of longitudinal recordings contained some missing values, they were analyzed by fitting a mixed model rather than by repeated measures ANOVA (which cannot perform analysis on data with missing values) (Prism 8.4.1, GraphPad).

**Figure 2.**
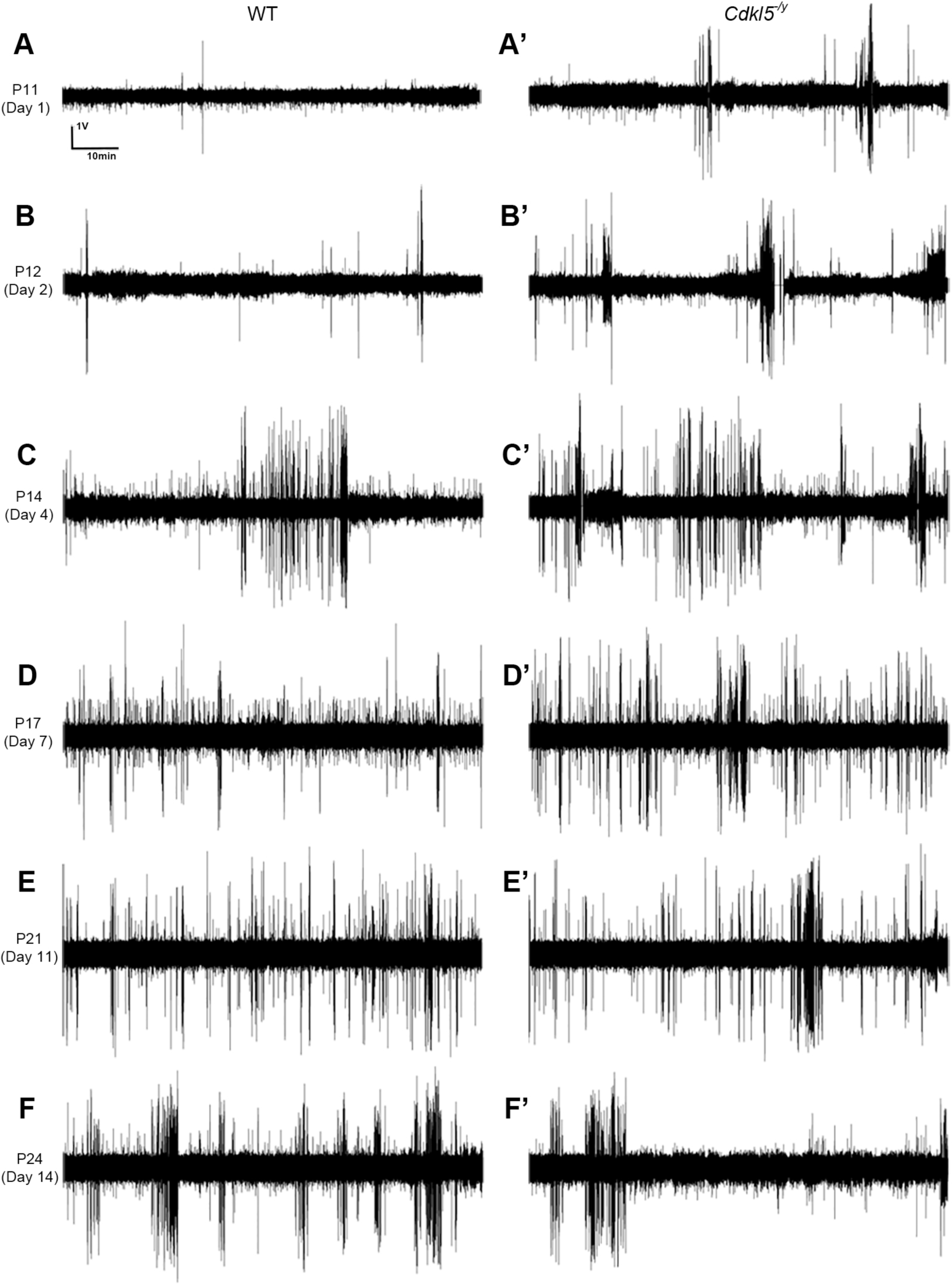
Longitudinal EEG activities recorded from mouse pups. Representative EEG activities of a pairs of *Cdkl5*^*-/y*^ and wild-type (WT) pups recorded for 100 minutes daily at P11, 12, 14, 17, 21, and 24. All the traces present the last 90 min (10 - 100 min). *Cdkl5*^*-/y*^ pups (A’-F’) display increased seizure-like discharges compared to WT pups (A-F) on the first three post-surgery days (A-C’); however, the number of bursts seems reduced in mutant compared to WT pup after weaning (P24, F and F’).

**Figure 3.**
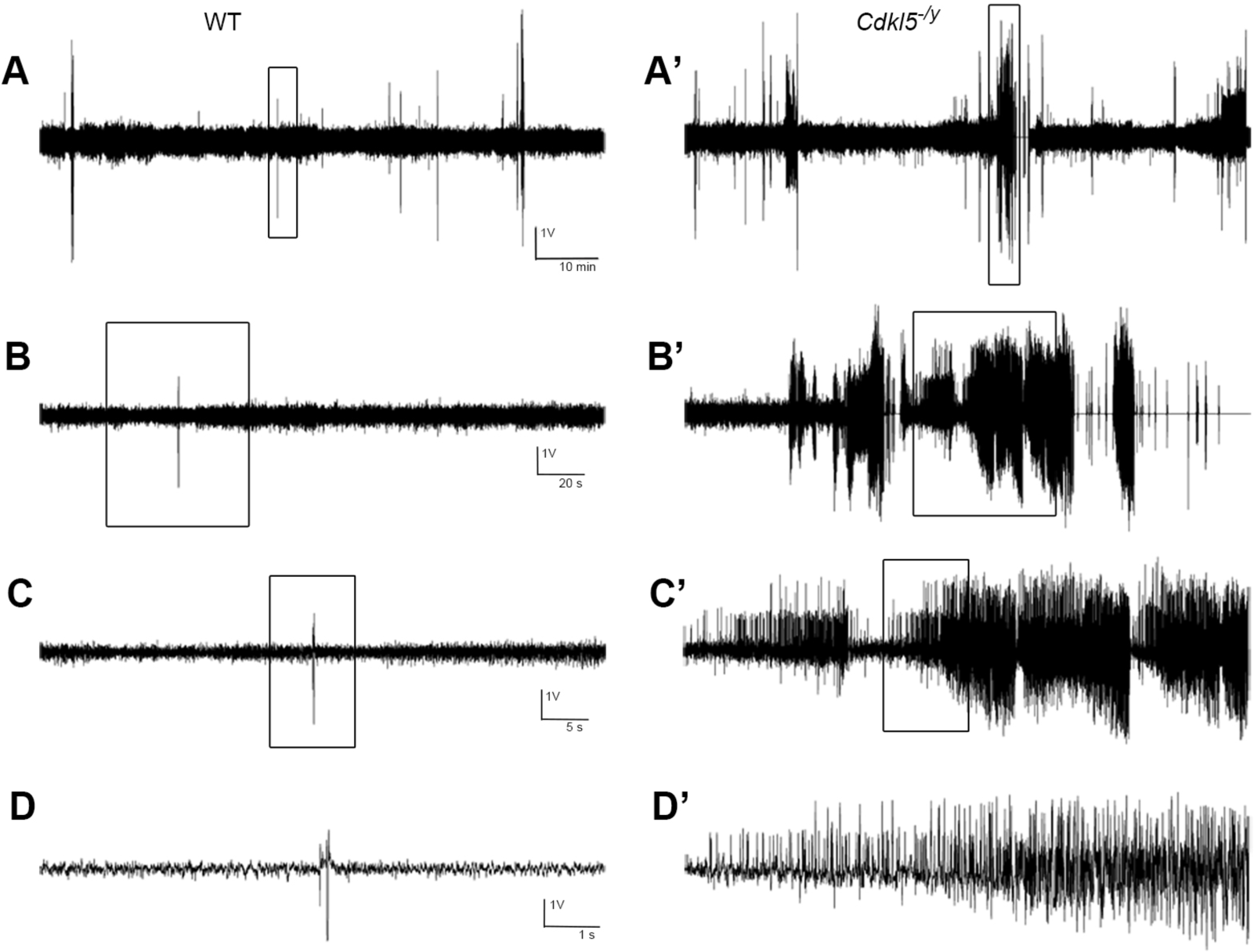
Seizure-like EEG activities found in *Cdkl5*^*-/y*^ pups at P12. Representative traces of EEG activities recorded from a pair of pups on the second day after implantation of transmitters. The trace in B (B’) is a zoomed view of the boxed area (4 min) in A. The boxed areas in B (B’, 60 sec) and C (C’, 10 sec) are magnified as shown in C (C’) and D (D’), respectively. Notably, the recurrent episodes of epileptiform discharges are manifested only in *Cdkl5*^*-/y*^ pups (A’-D’).

**Figure 4.**
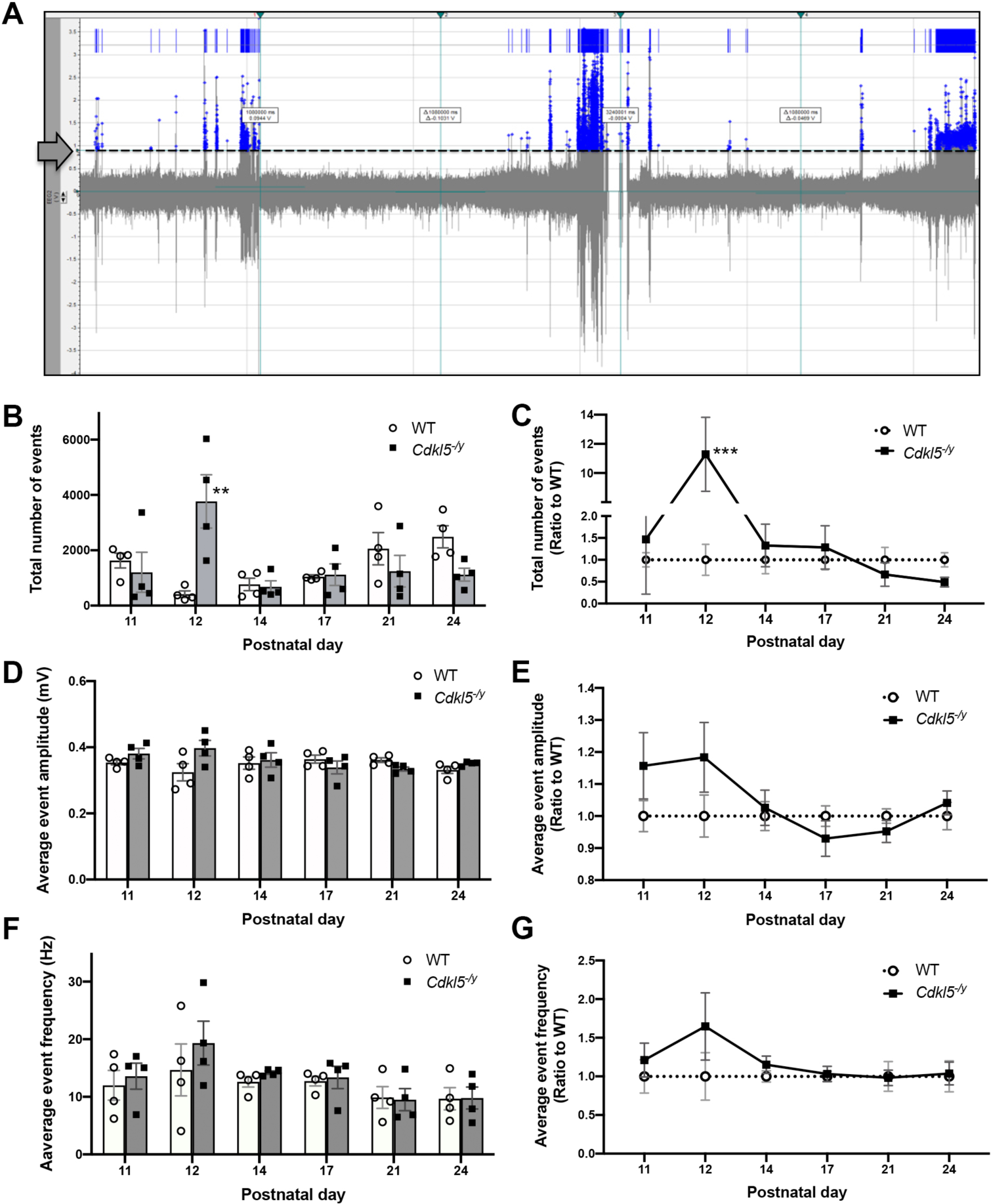
The number of events on EEG increased in *Cdkl5*^*-/y*^ pups at P12. Population spike discharges with amplitudes higher than 0.9 volt (set as the threshold, equivalent to 0.225 mV of input amplitude; arrow and dash line in A) were defined as “events” using the Clampex software. The blue points and the vertical bars on the top indicate the detected events. The total number of events is significantly increased in pups at P12, reaching to ∼12-fold of WT pups (B, C). Although *Cdkl5*^*-/y*^ pups show slight age-dependent alterations in the event amplitude (D, E) and frequency (F, G), the increase at P12 is not significant. All data are presented as mean ± SEM. ** *p* < 0.01; *** *p* < 0.001; mixed effects model followed by multiple comparison with Bonferroni’s test.

### Assessment for seizure-like behaviors

During the EEG recording, potential seizure-like behaviors were videotaped simultaneously using a webcam (Logitech). The pups’ behaviors at P12 selectively derived from the prolonged bursts whose durations were longer than the mean burst duration were evaluated according to a previous report (Price et al., 2009). The behavioral seizures were scored from 0 to 3 with the following criteria: 0, sleeping/ staying quiet without movement; 1, taking rest with occasional orofacial movement/ head bobbing/ forelimb twitch/ tail flick; 2, freezing suddenly during movement; and 3, spasm-like jerky movements of the head, body or limbs. The summed scores for spasm-like behaviors during the prolonged bursts for each *Cdkl5*^*-/y*^ pup (see Movie#6-1) were compared to their WT littermates (see Movie#6-2).

### Wire suspension

To test grip strength and endurance, the forepaws of male pups were positioned on a horizontal wire (3 mm in diameter) suspended at a height of 30 cm above the tabletop (Schneider and Przewlocki, 2005). Soft bedding materials were placed under the wire to protect fallen pups from injury. Each pup was tested for five trials every other day from P10 to P18. The latency to fall was recorded with a cutoff time of 3 min for each trial. The median of daily scores for *Cdkl5*^*-/y*^ pup (n = 11) and their WT littermates (n = 13) were compared by two-way repeated measures ANOVA (Prism 8.4.1, GraphPad).

### Grip strength measurement

Male mice (including the transmitter-implanted ones) at the age of 1-1.5 months were measured by a grip strength meter (UGO Basile, Italy); some of which were measured again at 5-7 months of age. The force transducer was placed at a height of ∼2.5 cm from the base plate. The tested mice were pulled against the T-shaped bar for 10 times with an inter-trial interval of 30 seconds to avoid muscle fatigue. The “peak force” (in grams, gF), “peak time” (i.e., the latency time of the animal response), and “total holding duration” (in seconds) were recorded. The rate of applied pulling force (target force rate, slope, i.e., gF/sec) was calculated to confirm the steady force of application. The “grip force per unit weight” (gF/g) was estimated from the mean of 10 trials of each mouse. Differences between genotypes were compared with mixed effects model followed by Bonferroni’s test.

### Open field test

The implanted *Cdkl5*^*-/y*^ mice (n = 5) and their littermate controls (n = 6) at P31 were tested as described previously (Su et al., 2015). Mice were videotaped in a transparent Plexiglas-made open field arena (40 × 40 × 25 cm) for 16 min with dim light. The total traveled distance of mice was analyzed for the last 12 min (excluding the first 3.5 min for habituation) with the Smart video tracking system (Harvard, USA). The detection threshold for body movements of locomotion was the speed of 2 cm per second. Differences between genotypes were compared by Student’s *t-*test.

### Immunoblotting

Immunoblotting was performed as previously described (Jhang et al., 2017). The cortical and hippocampal tissues were harvested from *Cdkl5*^*-/y*^ pups and their WT littermates (n = 3∼6) at the age of P12, and then homogenized by sonication in lysis buffer containing 1% protease inhibitors and 1% phosphatase inhibitor (Sigma). The protein lysates containing 20 micrograms of protein were separated in 4-15% polyacrylamide gel (Bio-Rad) with 125 V for 100 min. The proteins were transferred to PVDF membranes (Millipore) with 350 mA for 2 h (Mini Trans-Blot Cell, Bio-Rad). The membranes were blocked using 5% skim milk for 1 h and then incubated with primary antibodies against CDKL5 (1:1,000, Abcam), K^+^/Cl^-^ co-transporter 2 (KCC2; 1:1,000, Millipore), phosphorylated KCC2 at serine 940 (pKCC2-S940; 1:5,000, PhosphoSolutions), phosphorylated KCC2 at threonine 1007 (pKCC2-T1007; 1:5,000, PhosphoSolutions), Na^+^/K^+^/Cl^-^ co-transporter 1 (NKCC1; 1:10,000, Millipore), postsynaptic density protein 95 (PSD-95; 1:2,000, ThermoFisher), GABA_A_R-gamma 2 (GABA_A_R; 1:2,000, Millipore), glutamate decarboxylase 2 (GAD2; 1:1,000, Cell Signaling), cannabinoid receptor 1 (CB1R; 1:5,000, Abcam), vesicular GABA transporter (VGAT; 1:1,000, SYSY), vesicular glutamate transporter 2 (VGLUT2; 1:1,000, SYSY) and glyceraldehyde 3-phosphate dehydrogenase (GAPDH, 1:50,000, Millipore) at 4 °C for 16 h. Following incubation with peroxidase-conjugated secondary antibodies at room temperature for 2 h, the target proteins were detected by an enhanced chemoluminescence reagent (ECL, Millipore) with an image acquisition system (Biostep, Germany). After the pKCC2 were detected, the membranes were stripped and re-blocking, followed by incubation of primary antibodies for KCC2. The intensity of target protein expression was quantified with densitometry (Image J, NIH) and normalized with the loading control of GAPDH. The protein expression in mutants was presented as “ratio to WT” and the Student’*s t*-test was used for comparison between genotypes.

## Results

### Wireless EEG recording in mouse pups

To identify potential EEG abnormalities in *Cdkl5* null (*Cdkl5*^*-/y*^) mice, we implanted a transmitter on a mouse pup at P10, the youngest age possible for this experiment, and recorded EEG activity using a miniature telemetry system (Fig. 1). Occasionally, transmitters were damaged in the following days (2 in 12), likely gnawed by the dam. These data were excluded from analysis. Compared with the WT littermate controls, we found abnormal seizure-like discharges in *Cdkl5*^*-/y*^ pups starting from P11, one day after the transmitter was implanted (Fig. 1C, Fig. 2).

### Increased seizure-like discharges in Cdkl5^-/y^ pups

Given that the battery power of the transmitters has a fourteen-day guarantee, we collected EEG data from P11 to P24 upon implantation at P10. The results demonstrated that the population spikes tended to increase in WT with age (Fig. 2). By contrast, loss of CDKL5 apparently increased EEG activity in pups during the P11-14 period (Fig. 2A-C’), whereas reduced discharges at P24 compared to WT pups (Fig. 2F, F’). The latter observation was consistent with previous reports that adult *Cdkl5* null mice lack spontaneous seizures (Wang et al., 2012; Amendola et al., 2014; Okuda et al., 2017). Further analysis of the EEG activities at P12 revealed that epileptiform discharges occurred in *Cdkl5*^*-/y*^ pups in a repetitive manner with episodes lasting from 10 to 20 sec (Fig. 3B’, C’), which were not detected in WT littermate control pups (Fig. 3B, C).

### Loss of CDKL5 triggers spontaneous seizures at P12

To quantify the electrographic spikes, we processed the EEG traces of the last 90 min in each day of the recording (i.e., 10-100 min) using Clampex software. For event detection, the “threshold” amplitude was set to 0.9 volts based on our pilot tests (arrow and dash line in Fig. 4A). We found that the total number of events significantly differ between genotypes with age [interaction of genotype x age: *p* = 0.0003, F (5, 36) = 6.265], in which it was significantly higher in *Cdkl5*^*-/y*^ pups compared to that in WT pups at P12 (3766 ± 961.7 in KO vs. 410.3 ± 118.6 in WT, *p* < 0.01; Fig. 4B, C; Table 4-1). To compare the amplitude of spike events, we normalized the output spikes of 1V to 0.25 mV based on the amplification of the transmitters (4000X). We found that the mean event amplitude was indeed comparable between genotypes with age, although there was a trend of increase at P11 and P12 (Fig. 4D, E; Table 4-1). In addition, there was no significant alteration in the mean frequency of total events in *Cdkl5*^*-/y*^ pups compared to WT pups with age, except a slight increase found in *Cdkl5*^*-/y*^ pups at P12 (Fig. 4F, G; Table 4-1).

We next evaluated the seizure-like activities through a burst analysis, in which a “burst” was defined as a series of continuous discharges for more than 20 events whose inter-event-intervals were less than 1 sec. The “total number of bursts” was increased in *Cdkl5*^*-/y*^ compared to WT pups at P12, however, the burst number became comparable between genotypes from P14 to P24, demonstrating possible normalization of EEG activities in *Cdkl5*^*-/y*^ pups with age (Fig. 5A, B; Table 4-1). Notably, the “duration of bursts” and the “number of events per burst”, both indicating the severity of epileptic discharges, were significantly increased in *Cdkl5*^*-/y*^ pups compared to WT littermates at P12 (*p* < 0.001; Fig. 5C-F; Table 4-1). This enhancement appeared to be subsided and returned to WT levels during the following days. The frequency of events in burst, though, did not change in CDKL5-deficient pups compared to WT littermates through P11 to P24 (Fig. 5G, H; Table 4-1).

**Figure 5.**
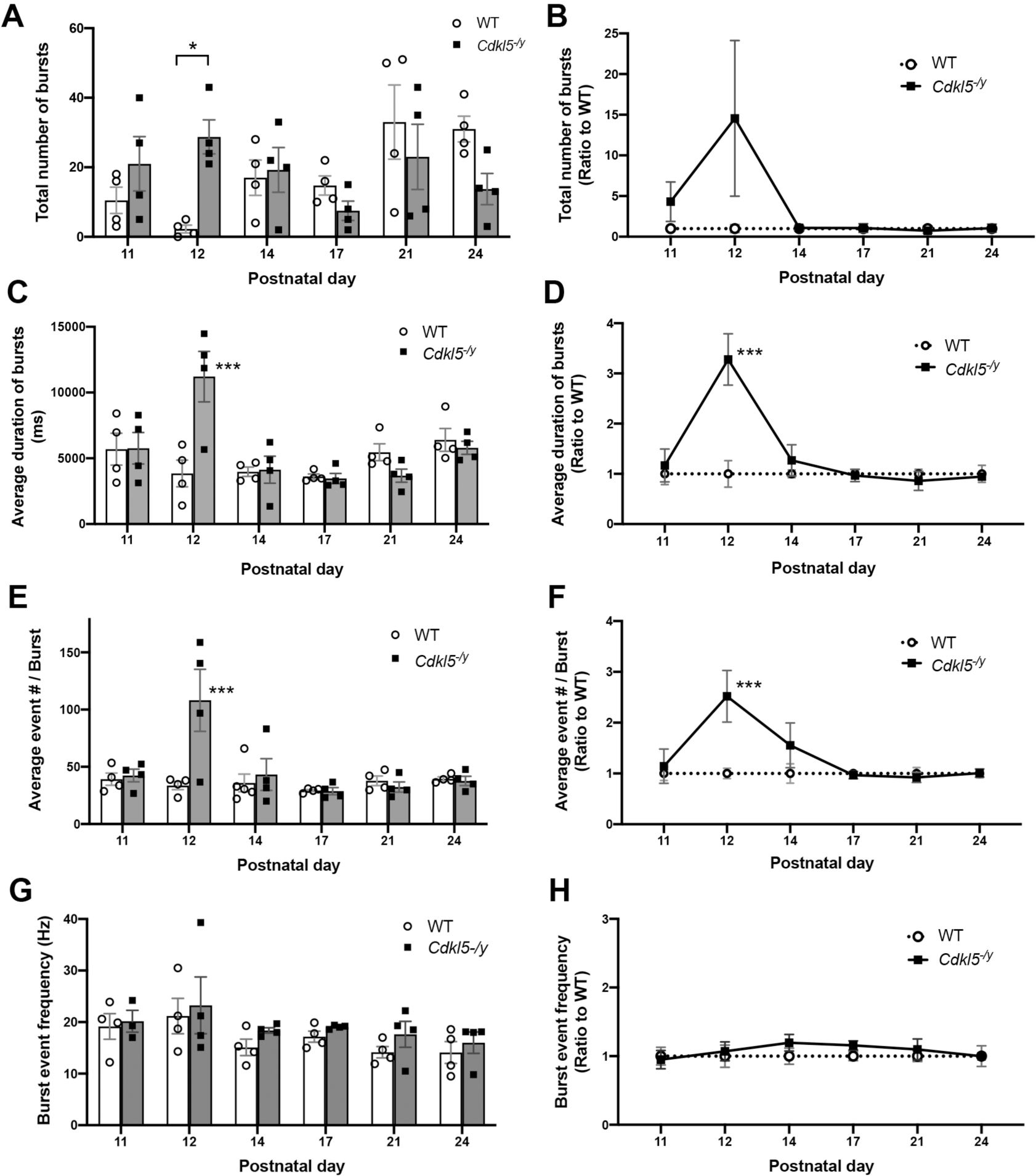
Increased burst activities on EEG detected in *Cdkl5*^*-/y*^ pups at P12. A burst was defined as > 20 continuous electrographic discharge events with the inter-event-interval for < 1 sec. *Cdkl5*^*-/y*^ pups exhibit an increased total number of bursts (A, B), duration of bursts (C, D) and event number per burst (E, F) compared to WT littermate pups. The mean frequency of bursting events (G, H), however, is comparable between genotypes. All data are presented as mean ± SEM. * *p* < 0.05; *** *p* < 0.001; mixed effects model followed by multiple comparison with Bonferroni’s test.

### Increased spasm-like movements in Cdkl5^-/y^ pups during the prolonged bursts

Given that behavioral seizures have been observed as a primary symptom of infantile epilepsy (Aujla et al., 2009; Marsh et al., 2009; Price et al., 2009), we next analyzed the videos captured simultaneously with EEG recordings to identify potential behavioral seizures that occurred during the electrographic events of bursts. We selectively reviewed the behaviors of pups at P12 that happened during the prolonged burst events, finding that these events were associated with frequent limb twitches, rapid limb stretching, tail flicks and occasional jerky movements of the whole body, in *Cdkl5*^*-/y*^ pups (Fig. 6A, upper panels; Movie#6-1). However, WT littermate pups tended to rest quietly without obvious movements during the bursting period (lower panels in Fig. 6A; Movie#6-2). Following quantification of these spasm-like behaviors by scoring them from 0 to 3, we found that the accumulative scores were consistently higher in *Cdkl5*^*-/y*^ pups compared to the scores derived from the WT littermate pups (Fig. 6B).

**Figure 6.**
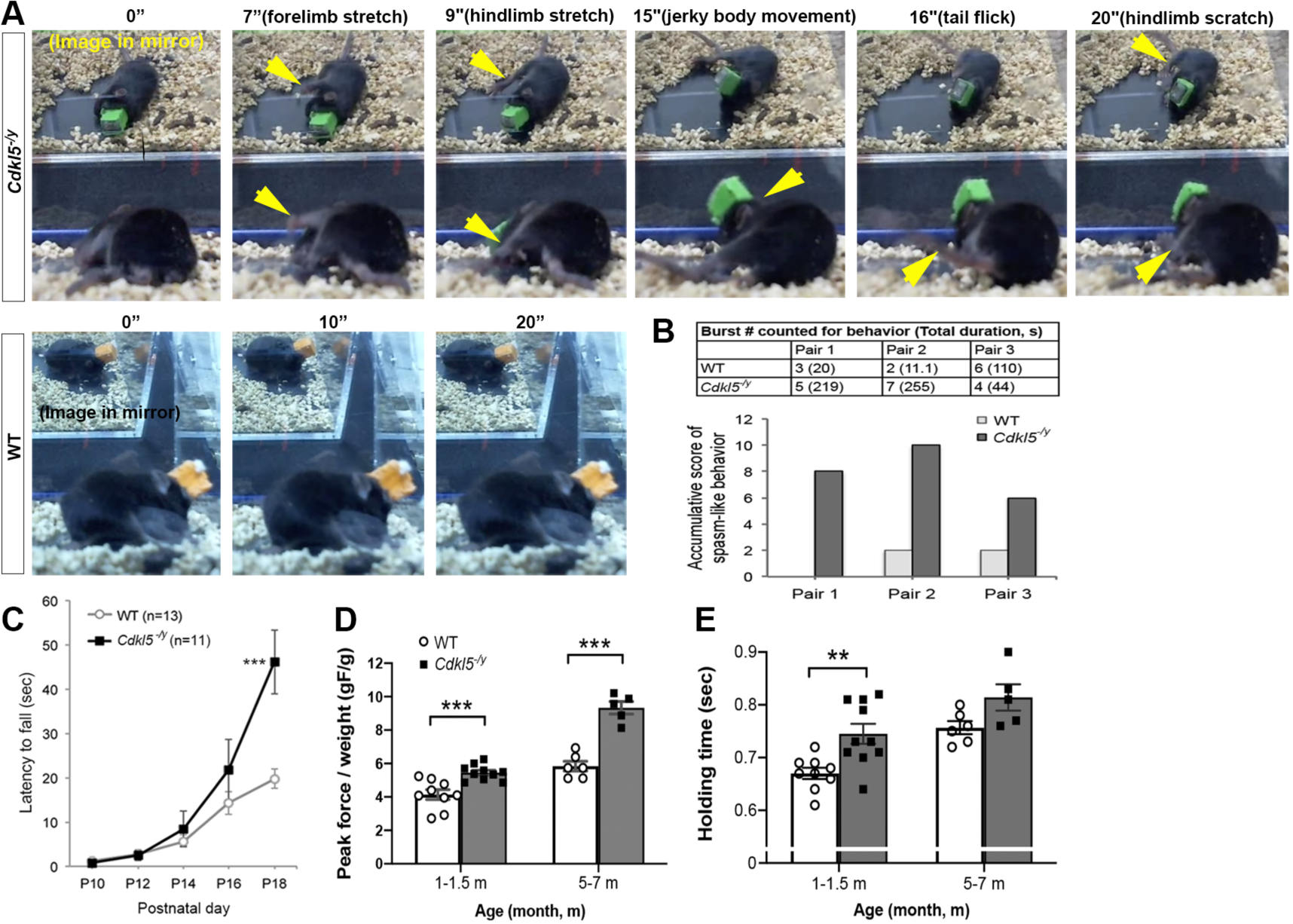
Spontaneous ictal behaviors developed in *Cdkl5*^*-/y*^ pups and adults. Representative screenshots from a video of *Cdkl5*^*-/y*^ pup shows continuous spasm-like movements within a prolonged burst of EEG activity (A, upper panel; also see Movie #6-1), while WT pup maintains the same posture throughout the burst of 20 sec (A, lower panel; also see Movie #6-2). The accumulative behavioral scores of spasm-like movements during the prolonged bursts were estimated in three pairs of pups (B). *Cdkl5*^*- /y*^ pups gradually developed ictal grasping prior to weaning (C) and increased grip strength in adulthood (D, E). All data are presented as mean ± SEM. ** *p* < 0.01; *** *p* < 0.001; Two-way repeated measures ANOVA followed by Bonferroni’s test in C; mixed effects model followed by Bonferroni’s test in D and E.

### Enhanced ictal grasping in Cdkl5^-/y^ mice

Given that “ictal grasping” has been identified in patients with frontal hyperkinetic seizures, which presents increased grasping forces after the seizure onset (Gardella et al., 2006; Leiguarda et al., 2008), we next measured the grasping strength in mouse pups from P10 to P18 of age using the wire suspension test. We found that *Cdkl5* mutant pups tended to grasp the wire longer than their WT littermate controls from P14, and the increased holding time become prominent at P18 (46.1 ± 7.2 sec in KO vs. 19.8 ± 2.1 sec in WT, *p* < 0.001; Fig. 6C). These results suggest that, in addition to triggering the onset of seizure-type EEG activities at P12, loss of CDKL5 may enhance ictal grasping later in life.

To clarify if the enhancement of ictal grasping persists to adulthood, we next examined grip strength of *Cdkl5*^*-/y*^ and WT mice by pulling their tails with a steady force at the age of 4-6 weeks. We found that the rate of pulling force (i.e., slope of peak force) was comparable between groups, suggesting that the force we applied was even and steady (212.04 ± 8.27 in KO vs. 190.59 ± 14.55 in WT, *p* > 0.05). As pre-weaning increase of grasping, *Cdkl5*^*-/y*^ juveniles exhibited increased grip strength (peak force, gF) upon normalization to body weight (gF/g: 5.45 ± 0.15 in KO vs. 4.14 ± 0.29 in WT, *p* < 0.001; Fig. 6D) and prolonged holding time of grasping compared to their littermate controls (0.74 ± 0.02 sec in KO vs. 0.67 ± 0.01 sec in WT, *p* < 0.01; Fig. 6E). These traits persisted with age, as similar grip strengthening was found in mutant mice at 5-7 months of age (gF/g: 9.33 ± 0.37 in KO vs. 5.84 ± 0.29 in WT, *p* < 0.001; Fig. 6D). Of note, we found that *Cdkl5*^*-/y*^ pups implanted with transmitters exhibited an increase of ∼1.8-fold in locomotor activities at P31 compared to their WT littermates in an open field test (Traveled distance: 5683.5 ± 450.7 cm in KO vs. 3222.1 ± 206.4 cm in WT, *p* < 0.001), suggesting that the neonatal implantation of transmitters and the longitudinal EEG recordings during the early postnatal age do not interfere the presentation of behavioral phenotypes, such as hyperlocomotion, in adulthood (Wang et al., 2012; Jhang et al., 2017).

### Reduced functional KCC2 in Cdkl5^-/y^ cortex at P12

CDKL5 has been reported to translocate into the nucleus by the age of P14 and may regulate gene expression through phosphorylation of transcriptional regulators (Rusconi et al., 2008). The developmental induction of the neuronal K^+^/Cl^-^ co-transporter 2 (KCC2), which extrudes intracellular chloride anions, has been reported to be essential for the ontogenetic switch of GABA_A_-mediated responses from depolarizing to hyperpolarizing (Rivera et al., 1999). The expression and function of KCC2 is thus a key to control neuronal excitability in neonates, and KCC2 dysfunction has been implicated in pathogenesis of infantile epilepsy (Puskarjov et al., 2014; Silayeva et al., 2015; Sivakumaran et al., 2015; Stodberg et al., 2015; Glykys et al., 2017; Moore et al., 2017). Notably, phosphorylation of KCC2 at the residue of serine 940 (pKCC2-S940) is crucial for stabilizing KCC2 on the surface of neuronal membrane that allows extrusion of Cl^-^ and maintains activity of KCC2 (Lee et al., 2007; Kahle et al., 2013). Given our findings of age-dependent epileptic activities in *Cdkl5*^*-/y*^ pups, it raises a possibility that CDKL5 deficiency may affect the level of protein expression and/or phosphorylation of KCC2. We therefore measured the phosphorylation and protein expression levels of KCC2 by immunoblotting the cortical protein lysates from mice carrying CDKL5 or not. We found that loss of CDKL5 significantly reduced the phosphorylation level of pKCC2-S940 as well as the proportion of pKCC2-S940 in total KCC2 compared to that in WT littermate controls (0.58 ± 0.056 of WT, *p* < 0.01; Fig. 7A, B). By contrast, phosphorylated KCC2 at threonine 1007 (pKCC2-T1007), which inhibits its transport function (Inoue et al., 2012), was not changed in *Cdkl5*^*-/y*^ pups (0.913 ± 0.071 of WT, *p* > 0.05; Fig. 7A, B). The alterations of phosphorylated KCC2 were recapitulated in the hippocampus of CDKL5-deficient pups at P12 (data not shown). Notably, we did not find a change in the expression of total KCC2 protein (1.005 ± 0.070 of WT, *p* > 0.05) and Na^+^/K^+^/Cl^-^ co-transporter 1 (NKCC1; 0.953 ± 0.041 of WT, *p* > 0.05) (Fig. 7A, C), a developmentally down-regulated cation-chloride co-transporter (Yamada et al., 2004).

**Figure 7.**
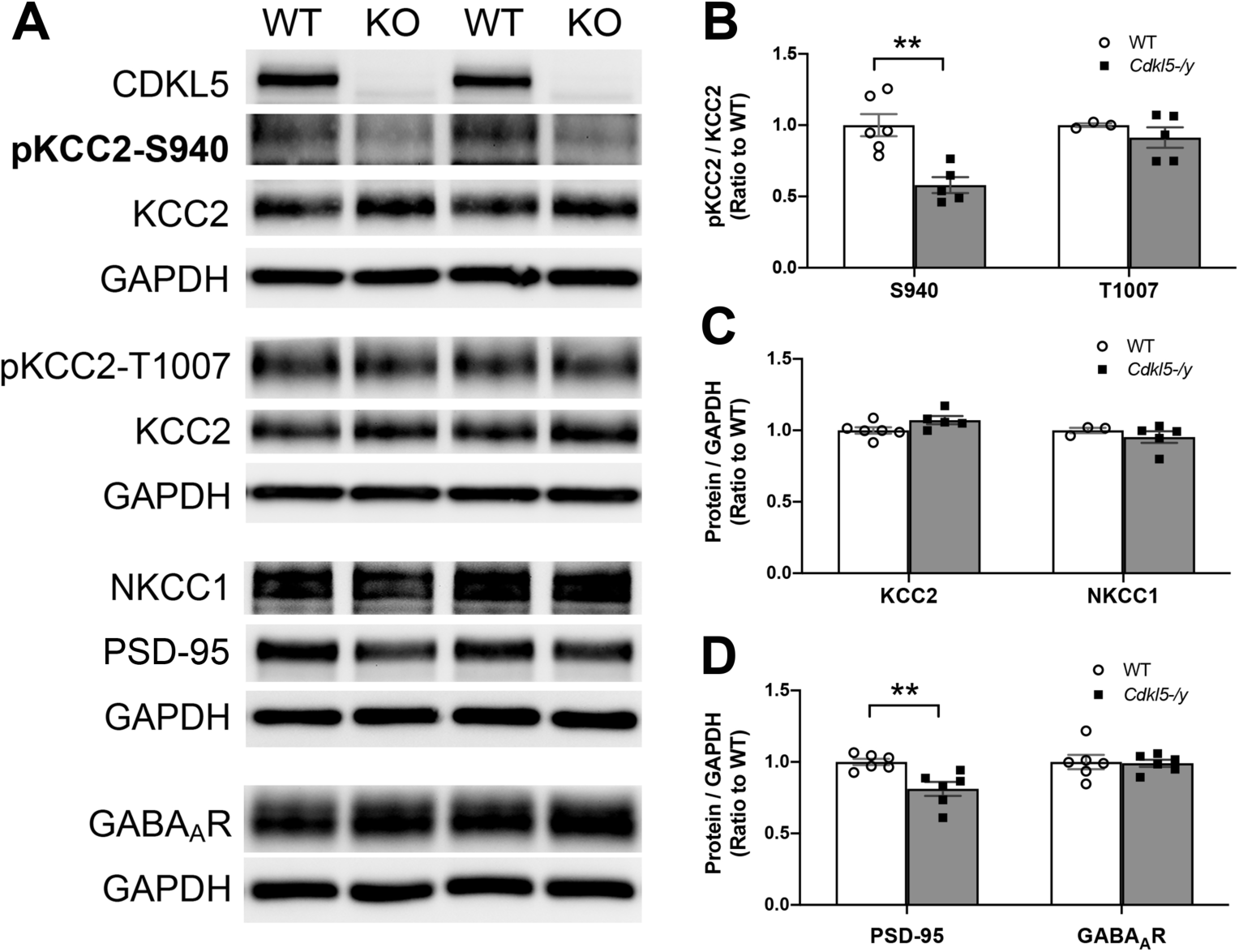
Phosphorylation of KCC2 reduced in the cortex of *Cdkl5*^*-/y*^ pups at P12. Immunoblotting shows that loss of CDKL5 reduces the level of phosphorylated KCC2 at serine 940 (pKCC2-S940; A, B) in the rostral cortex at P12. No significant alteration is found in protein expression of total KCC2, phosphorylated KCC2 at threonine 1007 (pKCC2-T1007), NKCC1 and GABA_A_ receptor. The reduction of PSD-95 in *Cdkl5*^*-/y*^ pups (A, D), consistent with the findings reported previously, serves as a positive control. The protein expression was quantified by densitometry with Image J using GAPDH as a loading control. The results are derived from three litters and presented as mean ± SEM. ** *p* < 0.01, Student’s *t-*test.

We further examined the expression level of candidate proteins related to neuronal inhibition, finding that GABA_A_ receptors (0.991 ± 0.025 of WT, *p* > 0.05; Fig. 7A, D), GABA synthesizing enzyme glutamate decarboxylase 2 (GAD2; 1.063 ± 0.011 of WT, *p* > 0.05; n = 3) and vesicular GABA transporter (vGAT; 1.017 ± 0.026 of WT, *p* > 0.05; n = 3) are evenly expressed between mutant and WT mice, at least in the cortical tissues. The expression levels of vesicular glutamate transporter 2 (vGluT2; 0.983 ± 0.027 of WT, *p* > 0.05; n = 3) and cannabinoid receptor 1 (CB1R; 0.937 ± 0.071 of WT, *p* > 0.05; n = 3), the excitatory presynaptic proteins, were not different between genotypes either. However, the protein level of PSD-95, a CDKL5-interacting protein in the postsynaptic density, appeared to be reduced in pups lacking CDKL5 (0.812 ± 0.049 of WT, *p* < 0.01; Fig. 7A, D), consistent with previous studies in mutant mice and in cultured neurons (Ricciardi et al., 2012; Zhu et al., 2013; Della Sala et al., 2016; Pizzo et al., 2016; Lupori et al., 2019). Thus, KCC2 phosphorylation at S940 appears to be selectively down-regulated in *Cdkl5* null pups, especially at the age when they are suffering from spontaneous seizures.

## Discussion

In the present study, we measured EEG in neonatal mice using wireless setup across different developmental time points from P11 to P24, and videotaped their behaviors in parallel. Upon quantification of EEG discharges, we found that pups with CDKL5 deficiency exhibited increased epileptiform discharges selectively at P12, with a drastic increase in the total number of spike events, the total number of bursts, the mean duration of bursts and the mean event number per burst compared to WT littermate controls. These mutant pups also showed seizure-like behaviors, such as limb stretch, tail flicks and jerky body movement, during the prolonged bursting period at P12. Enhanced ictal grasping, manifested in *Cdkl5* mutant pups at late pre-weaning stage, was sustained to adulthood with an increase in grip strength and holding duration. Moreover, loss of CDKL5 significantly decreased the phosphorylation level of pKCC2-S940 and reduced the proportion of functional KCC2 in the developing cortex. Our findings suggest that loss of CDKL5 drives epileptic activities in neonatal mice at P12, and this phenotype is likely the consequence of extended GABA-mediated excitation due to impaired function of KCC2.

As a mouse model of CDD, *Cdkl5*^*-/y*^ mice have exhibited numerous behavioral phenotypes recapitulating features in CDD (Wang et al., 2012; Amendola et al., 2014; Jhang et al., 2017; Mazziotti et al., 2017; Okuda et al., 2018). However, early-onset epilepsy, the symbolic trait in children with CDD, has not been reported in previous studies. Given that eighty percent of children with CDD have daily seizures, and the epileptic spasms onset at a median age of 6 weeks followed by a seizure-free honeymoon period (Fehr et al., 2016; Demarest et al., 2019; Olson et al., 2019), we reasoned that onset of epilepsy in mice with CDKL5 deficiency may occur during the early postnatal age and likely become alleviated later in life. By using a miniature telemetry system for wireless EEG recording in mouse pups, we identified robust epileptiform discharges in pups with CDKL5 deficiency selectively at P12. These epileptic EEG activities were not caused by contamination of muscular contractions because there was barely ictal behaviors (Fig. 6A) occurred prior to the start of prolonged bursts in *Cdkl5* null pups (data not shown). We noticed a trend of increase in total number of both events and bursts in WT pups from P12 to P24 (Fig. 4B and 5A). With increased activities in WT pups, EEG profiles in *Cdkl5*^*-/y*^ pups appeared to be slightly reduced compared to WT at P24 (Fig. 4C), which is consistent with previous findings of reduced number of epileptiform events in *Cdkl5*^*-/y*^ adult mice (Amendola et al., 2014). Interestingly, our findings of the transient epileptic EEG activities in CDKL5-deficient pups echoes a recent study that demonstrates an arrest of hyperexcitability in the early stage of developing cortical organoid derived from CDD patients (Negraes, 2019). The mechanism for the CDKL5-mediated transient control of neuronal excitability remains to be investigated. Our findings do not exclude the possibility that seizure onset occurs prior to P12 in CDKL5-deficient mice, which is barely examined due to the plight that pups younger than P10 are too vulnerable to survive from implantation of transmitters.

To elucidate possible mechanisms underlying the age-dependent early onset of seizures, we measured the expression of candidate proteins involved in the developmental switch of hyperpolarizing GABAergic inhibition, which has been implicated in pathogenesis of neonatal seizures (Rivera et al., 1999; Puskarjov et al., 2014; Silayeva et al., 2015; Sivakumaran et al., 2015; Stodberg et al., 2015; Glykys et al., 2017; Moore et al., 2017). We found that CDKL5 deficiency significantly decreased the cortical level of pKCC2-S940 by 42% at P12 (Fig. 7). The KCC2 protein does not express in rat hippocampus until P5 and reaches to the adult level by P15 (Rivera et al., 1999; Watanabe and Fukuda, 2015), corresponding to ∼P13 in mice (Clancy et al., 2001; Li et al., 2002). Our findings that loss of CDKL5 reduced the proportion of functional KCC2 (pKCC2-S940) without altering the expression of total KCC2 at P12 suggest that loss of CDKL5 may cause hypofunction of KCC2, leading to delayed transition of GABAergic response from excitation to inhibition and resulting in seizure onset in *Cdkl5*^*-/y*^ pups at the neonatal stage. Supporting of this notion, previous studies have found that onset of kainate-induced status epilepticus is coupled with S940 dephosphorylation and internalization of KCC2, and is accelerated by mutation of S940 (Silayeva et al., 2015). Recessive KCC2 mutations, decreased KCC2 surface expression and loss of KCC2 activity have been reported in infants with a severe early-onset epileptic encephalopathy (Stodberg et al., 2015). Moreover, reduction of KCC2 and delayed functional switch of GABA has been reported in neurons from Rett syndrome (RTT) patients, suggesting the possible existence of a common mechanism for the epileptic trait in CDD and RTT (Tang et al., 2016; Kadam et al., 2019). Therefore, CDKL5 may play a key role to preserve KCC2 function and modulate the initiation of GABAergic inhibition during neonatal stage. The intracellular location of KCC2, the characteristics of KCC2-positive neurons and the excitabilities of GABA receptive neurons in neonatal CDKL5-deficient mice remain further investigation in future studies.

On the other hand, given that KCC2 has not been identified as a direct substrate of CDKL5 (Baltussen et al., 2018; Munoz et al., 2018), the CDKL5-dependent regulation of pKCC2 might be mediated indirectly through the action of protein kinase C, which has been reported to interact with CDKL5 and phosphorylate KCC2 at S940 (Lee et al., 2007; Wang et al., 2012). Although we did not find any alteration in the expression of proteins related to excitation/inhibition (E/I) balance, such as pKCC2-T1007, NKCC1, GABAA receptor, GAD2, vGAT, vGluT2 and CB1R, in the cortex at P12, other players, such as NMDA receptors, AMPA receptors and parvalbumin, could still play a role in CDKL5-deficiency-induced E/I imbalance (Pizzo et al., 2016; Okuda et al., 2017; Tang et al., 2017; Yennawar et al., 2019).

Despite spontaneous grasping of objects in many clinical cases of CDD at a few months of age (Bartnik et al., 2011), the majority of children with CDD exhibit delayed raking grasp, especially in males (Fehr et al., 2015). Our findings of persistent enhancement in grasping duration and strength in both neonatal and adult mice (Fig. 6) suggest that the grasping behavior could be a robust trait representing an outcome measure for the hypermotor-tonic-spasms type of epilepsy in the frontocentral region (Gardella et al., 2006; Leiguarda et al., 2008; Klein et al., 2011; Melikishvili et al., 2019). In view of the expression of KCC2 in human, it reaches to the equivalent level as that in P10-15 rat at the early infancy stage (Kang et al., 2011), corresponding to the period of seizure occurrence in children with CDD (Fehr et al., 2016; Demarest et al., 2019). Our findings of epileptic discharges in *Cdkl5*^*-/y*^ mouse pups at P12, therefore could be temporally relevant to the time window of seizure onset in children with CDD, in terms of KCC2 expression. Given that children with CDD are often not responsive to AEDs or responding without long-term efficacy (Muller et al., 2016; Dale et al., 2019; Olson et al., 2019), the KCC2 activators could be a potential alternatives for CDD treatment based on our findings.

Together, in the present study, we identified early-onset seizures, from both EEG and video recordings, in neonatal CDKL5-deficient mice, offering construct and face validity for this mouse model of CDD. We also demonstrated that CDKL5 is required for regulation of KCC2 phosphorylation, providing a potential diagnostic biomarker and therapeutic target for CDD. By leveraging gestational diagnosis and intervention, early onset seizures and the consequent developmental delays in children with CDD would be possibly overcome in the near future.

## Acknowledgments

We thank Dr. Jin-Chung Chen for providing equipment for grip strength measurement and critical reading of the paper. This work is supported by grants from the Ministry of Science and Technology, Taiwan [MOST107-2320-B-004-001-MY3 to WL; MOST108-2636-B-110-001 to KZL] and the International Foundation of CDKL5 Research, USA [2015∼2019 to WL].

## Extended data

Table 4-1. Quantification and statistic results for longitudinal measurements of EEG discharges from mouse pups

Movie#6-1. Representative behavior of a *Cdkl5*^*-/y*^ pup during a prolonged burst at P12

Movie#6-2. Representative behavior of a WT pup during a prolonged burst at P12

