## Supplementary material for "Deficiency of cyclin-dependent kinase-like 5 causes spontaneous epileptic seizures in neonatal mice": Table 4-1

**Table 4-1. Quantification and statistic results for longitudinal measurements of EEG discharges from mouse pups**

| Mean ± SEM, (n = 4) | P11 | P12 | P14 | P17 | P21 | P24 |
|---|---|---|---|---|---|---|
| Total number of events (Fig. 4B) | WT: 1627.3 ± 265.4<br>KO: 1205.0 ± 724.7<br>$p > 0.05$ | WT: 410.3 ± 118.6<br>KO: 3766 ± 961.7<br>$p < 0.001$ (***) | WT: 766.8 ± 224.3<br>KO: 680.3 ± 214.3<br>$p > 0.05$ | WT: 1044.8 ± 67.7<br>KO: 1119.5 ± 390.1<br>$p > 0.05$ | WT: 2058.0 ± 581.5<br>KO: 1243.8 ± 569.7<br>$p > 0.05$ | WT: 2489 ± 399.7<br>KO: 1117.3 ± 232.5<br>$p > 0.05$ |
| Average event amplitude (mV) (Fig. 4D) | WT: 0.354 ± 0.007<br>KO: 0.381 ± 0.016<br>$p > 0.05$ | WT: 0.324 ± 0.026<br>KO: 0.397 ± 0.024<br>$p > 0.05$ | WT: 0.352 ± 0.019<br>KO: 0.362 ± 0.022<br>$p > 0.05$ | WT: 0.364 ± 0.012<br>KO: 0.339 ± 0.020<br>$p > 0.05$ | WT: 0.361 ± 0.007<br>KO: 0.334 ± 0.006<br>$p > 0.05$ | WT: 0.331 ± 0.011<br>KO: 0.351 ± 0.003<br>$p > 0.05$ |
| Average event frequency (Hz) (Fig. 4F) | WT: 12.0 ± 2.6<br>KO: 13.6 ± 2.3<br>$p > 0.05$ | WT: 14.7 ± 4.5<br>KO: 19.3 ± 3.8<br>$p > 0.05$ | WT: 12.6 ± 0.9<br>KO: 14.2 ± 0.3<br>$p > 0.05$ | WT: 12.7 ± 0.8<br>KO: 13.4 ± 1.9<br>$p > 0.05$ | WT: 9.9 ± 1.9<br>KO: 9.5 ± 1.9<br>$p > 0.05$ | WT: 9.7 ± 1.9<br>KO: 9.8 ± 1.9<br>$p > 0.05$ |
| Total number of bursts (Fig. 5A) | WT: 10.5 ± 3.8<br>KO: 21 ± 7.8<br>$p > 0.05$ | WT: 2.3 ± 1.1<br>KO: 28.8 ± 4.9<br>$p < 0.05$ (*) | WT: 17.0 ± 5.1<br>KO: 19.3 ± 6.4<br>$p > 0.05$ | WT: 14.8 ± 2.8<br>KO: 7.5 ± 2.8<br>$p > 0.05$ | WT: 33.0 ± 10.7<br>KO: 23.0 ± 9.4<br>$p > 0.05$ | WT: 31.0 ± 3.7<br>KO: 13.8 ± 4.5<br>$p > 0.05$ |
| Average duration of bursts (ms) (Fig. 5C) | WT: 5692 ± 1210.2<br>KO: 4560.5 ± 1311.0<br>$p > 0.05$ | WT: 3848.4 ± 1013.4<br>KO: 11211.7 ±1923.5<br>$p < 0.001$ (***) | WT: 3980 ± 350.4<br>KO: 4129.3 ± 1027.2<br>$p > 0.05$ | WT: 3611.2 ± 193.1<br>KO: 3463.3 ± 389.7<br>$p > 0.05$ | WT: 5461.3 ± 640.7<br>KO: 3679.8 ± 496.9<br>$p > 0.05$ | WT: 6397.5 ± 233.4<br>KO: 5792.2 ± 498.1<br>$p > 0.05$ |
| Average event number per burst (Fig. 5E) | WT: 39.2 ± 5.4<br>KO: 42.4 ± 5.5<br>$p > 0.05$ | WT: 36.3 ± 4.8<br>KO: 108.1 ± 27.1<br>$p < 0.001$ (***) | WT: 37.8 ± 9.7<br>KO: 43.2 ± 13.8<br>$p > 0.05$ | WT: 29.3 ± 0.9<br>KO: 28.7 ± 3.1<br>$p > 0.05$ | WT: 37.8 ± 4.2<br>KO: 32.3 ± 4.5<br>$p > 0.05$ | WT: 39.5 ± 1.7<br>KO: 37.7 ± 4.1<br>$p > 0.05$ |
| Burst event frequency (Hz) (Fig. 5G) | WT: 19.2 ± 2.5<br>KO: 20.1 ± 1.8<br>$p > 0.05$ | WT: 21.2 ± 3.4<br>KO: 23.3 ± 5.5<br>$p > 0.05$ | WT: 15.1 ± 1.6<br>KO: 18.4 ± 0.5<br>$p > 0.05$ | WT: 17.2 ± 1.1<br>KO: 19.0 ± 0.2<br>$p > 0.05$ | WT: 14.2 ± 1.1<br>KO: 17.6 ± 2.5<br>$p > 0.05$ | WT: 14.1 ± 2.1<br>KO: 16.0 ± 2.1<br>$p > 0.05$ |
